# A mathematical model for storage and recovery of motor actions in the spinal cord

**DOI:** 10.1101/2020.05.27.119321

**Authors:** David J Parker, Vipin Srivastava

**Affiliations:** Department of Physiology, Development and Neuroscience, University of Cambridge, Cambridge CB2 3EJ, UK; School of Physics, University of Hyderabad, Hyderabad 500046, India

**Keywords:** motor learning, locomotion, orthogonalisation, normalisation

## Abstract

Motor outputs are generated by the spinal cord in response to descending inputs from the brain. While particular descending commands generate specific outputs, how descending inputs interact with spinal cord circuitry to generate these outputs remains unclear. Here, we suggest that during development particular motor programmes are stored in premotor spinal circuitry, and that these can subsequently be retrieved when the associated descending input is received. We propose that different motor patterns are not stored in the spinal cord as a library of separate programmes, but that the spinal cord orthogonalises and normalises the various inputs, identifies the similarities and differences between them, and stores only the differences: similarities between patterns are recognised and used as a common basis that subsequent input patterns are built upon. By removing redundancy this can greatly increase the storage capacity of a system composed of a finite number of processing units, thus overcoming the problems associated with the storage limits of conventional artificial networks (e.g. ‘catastrophic interference’). Where possible we relate the various stages of the processing to the known circuitry and synaptic properties of spinal cord locomotor networks, and suggest experimental approaches that could test unknown aspects.

## 1. Introduction

It is generally accepted that spinal cord networks can generate basic motor outputs in response to chemical or electrical stimulation independently of any descending or sensory inputs (i.e. ‘fictive’ locomotion; Kiehn et al. 2010). Spinal cord locomotor networks in isolation have been studied in the context of relatively simpler rhythmic motor outputs (e.g. swimming, flying, walking). These behaviours can take various forms (e.g. alternating left/right or flexion/extension at different frequencies, or struggling or synchronous activity), reflecting differences in the type of muscles recruited, and the timing and magnitude of their activation (Bizzi et al. 2008). From fictive locomotor studies it is assumed that the degree of excitation evoked by pharmacological or electrical stimulation determines the frequency of the motor outputs (Brodin et al 1985; Talpalar and Kiehn 2010; Cazalets et al 1992; Li and Moult 2012). However, the output is not directly proportional to the excitatory drive as stronger stimulation can cause qualitative changes in motor outputs (e.g. a switch from swimming to struggling movement; Roberts et al. 2010), and the effects of any stimulation are variable both within and between studies (see Cowley et al 2005). The latter property may reflect the extent to which normal locomotor rhythms are generated by the isolated spinal cord: pharmacologically-evoked fictive outputs occur in the absence of normal system interactions with sensory and descending inputs; reflects the tonic activation of all spinal cord neurons, and thus does not reflect natural activity; and it takes time to develop to a stable rhythm and has a variable success rate (see Li et al. 2006; Soffe et al. 2009; Ayers et al. 1983; Jackson et al 2005; Wang and Jung 2002; see Parker and Srivastava 2013 for discussion).

While fictive activity shows that the spinal cord contains the machinery needed to generate basic motor outputs, goal-directed movements (for example a particular speed of forward locomotion; Ausborn et al. 2019) are triggered by inputs that descend from the brain into the spinal cord. Unpatterned descending inputs evoked by stimulating brainstem locomotor regions can be converted by the spinal cord into different forms of patterned motor output (Shik and Orlovsky 1976). There are various descending pathways each with different targets within the spinal cord and thus different functional roles (Lemon 2008). Deliagina et al. (2002) addressed the decoding of descending inputs in the lamprey, and showed that different bilateral or unilateral commands can generate swimming, left or right turns, or up and down movements of the body. That different inputs to the spinal cord can generate specific outputs is also evidenced by microstimulation experiments that activate groups of neurons in the spinal cord or motor cortex (Bizzi et al. 2008).

However, how descending commands are converted into motor outputs is still not well understood (see Ausborn et al 2019). One suggestion is that the selection of motor outputs by descending inputs reflects a modular arrangement where spinal cord interneurons are organized into functional groups that are responsible for generating the muscle activity needed to evoke a certain movement (Bizzi et al 2008; Flash and Bizzi 2016). During development, descending inputs could become hard-wired to specific modules to provide dedicated circuitry through which a specific input can evoke a specific motor output (Graziano 2006; Goulding 2009). However, these circuits could also be modified as a result of motor learning to allow them to store and generate different motor outputs that could be recruited by particular descending commands (Wolpaw and Tennissen 2001). These possibilities are, of course, not mutually exclusive.

The spinal cord has the ability to store the results of motor learning (Thompson 2009; Thompson and Wolpaw 2014). However, even for relatively simple responses the mechanisms underlying this learning, as in the rest of the CNS, are unclear, both in terms of the underlying cellular and synaptic mechanisms, and how these changes relate to the actual behaviours that need to be stored and retrieved (Ausborn et al 2019). Here we address the issue of how the spinal cord could store motor programmes that can subsequently be triggered by differing descending commands. We don’t address specific types of motor outputs underlying different types of behaviours, but provide a mathematical description of how the spinal cord could in principle economically store a range of motor outputs that could be triggered by specific descending commands. Where possible, we place the proposed mathematical scheme in terms of the known circuitry of the lamprey spinal cord locomotor network (Buchanan 2001; Grillner 2003) and identified synaptic properties (Parker 2006; Jia and Parker 2016). As a complete cellular description is not possible even for this simpler system (Parker 2010), we suggest how the proposed mechanism can be tested experimentally in spinal cord circuits.

## 2. Storage of motor actions

We propose that motor outputs triggered by specific types of descending input from the brain are not stored in the spinal cord in their entirety as separate motor programs each time a new behaviour is learnt. The spinal cord instead identifies the similarities and differences between two or more motor programs and only stores the differences: the similarities are recognised and used as a common basis that subsequent motor programs are built upon (Srivastava et al. 2008; Srivastava and Sampath 2018). Thus, the common features of motor memories/programs are shared, and a specific motor program reflects the differences that distinguish it from other programs. This offers an economical solution by removing redundant parts of the signal, and can greatly increase the storage capacity of a system composed of a finite number of processing units by overcoming the ‘catastrophic interference’ (French 1999) that results when the same units are used to store different features (Srivastava and Edwards 2000; Srivastava et al. 2014). The spinal cord will contain a number of these stored patterns, *p*.

To start, assume that the nervous system is naive, for example at an early developmental stage, and that the spinal cord has no stored motor programs (though it could still respond reflexively to sensory inputs). A particular motor command will be relayed into the spinal cord through descending pathways (Graziano 2006; Lemon 2008). This descending command will reflect the activation of a subset of descending neurons that will become associated with a particular motor output. This input can be considered as a vector (see Schwartz 2016), whose components are the population of active and inactive cells (+1*/* −1, respectively), e.g.:

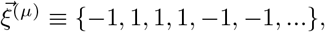

where the components +1*/* −1 are *N* in number; *N* being the number of spinal cord neurons receiving the input and the index *μ* identifies a particular input. Our hypothesis is that the *orthogonalised* versions of these vectors, to be denoted as 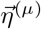, are stored in their *normalised* form, which we will denote as 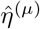. Vector normalisation is essential for treating all the vectors equally. This can be ensured if all the vectors are made to have the same length even though the individual components may differ from one another in a vector. Biologically this may mean that the net efficacy of each (normalised) vector is about the same. Also note that while the components +1*/* −1 in a 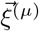 indicate whether the cells are active or inactive, the components in the corresponding 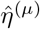 are fractions and are hypothesised to indicate the *degrees* of activity/inactivity of the cells. This may be considered as an added inherent virtue of ortho-normalisation, which seems more biologically appropriate than binary effects that only reflect spiking.

To describe how this is done in the lamprey spinal cord locomotor network, first consider the input, 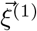. We hypothesise that this is received by a population of approximately 50 excitatory spinal cord interneurons (EIN; Buchanan 2001). Similar interneurons provide the excitation in other spinal cord networks (Ampatzis et al. 2014; Kiehn et al. 2010; Li et al. 2006, 2009), and are likely to be essential in the activation of spinal cord circuits. The EINs receive descending inputs to the spinal cord and provide excitatory inputs to motor neurons and other components of the locomotor network (Buchanan 2001). We suggest that in addition to relaying the descending input to motor neurons (Buchanan 2001), the EINs relay the input to a class of inhibitory interneurons (SiIN; see Buchanan 2001) to which they make monosynaptic connections (Parker 2003b). Inhibitory interneurons of this sort are again common components of spinal cord networks (Higashijima et al. 2004; Kiehn et al. 2010; Roberts et al. 2010). The descending input may project to both the EINs and SiINs. In fact we will invoke this except for the first input 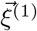 (for analytical reasons we assume that the system is initially a *tabula rasa*, so there is a ‘first’ input, which can be treated differently from those that follow it).

The EINs are functionally heterogeneous (Parker 2003a). This heterogeneity should influence the functional effects of the different EINs, and significantly in this context, it affects their potential for plasticity (Parker 2015). For the mechanism we propose here, we suggest that the EINs form three groups: group A receives descending inputs, sends outputs to EINs in group B (explained below), and receives feedback inhibitory inputs from the SiINs; group B doesn’t receive descending inputs from the brain but receives 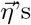 from A and transmits the normalised 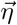, i.e. 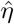 to the SiINs (note that group A EINs can receive inhibitory inputs from SiINs but can transmit signals to SiINs only via group B EINs); group C also does not receive the descending input but is connected with MNs. The EINs in group C receive the same inhibitory input from the SiINs as the group A EINs, but they do not send inputs to the SiINs. The different groups have different functional properties that we relate to experimentally identified features of these cells (Parker 2003a).

We suggest that a descending input for activating a particular motor output is stored in the dendrites of the pool of SiINs. Note that the orthogonalised version of the first input is the input itself, which is stored in the normalised form as 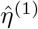. Normalisation has been suggested to be a canonical form of neural processing that computes the ratio between an individual neuron’s response and the summed activity of a group of neurons (Carandini and Heeger 2012). We suggest that the EINs in group B normalise the initial input relayed to them from group A and send the various components of vector 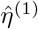, namely 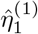, 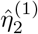, …, to the SiINs (we will detail normalisation below when we consider how a second descending command is stored).

These are then multiplied pair-wise on the dendrites of the SiINs (Koch and Poggio 1992) and lead to Hebbian-evoked changes in the EIN synapses made onto the SiINs in the following manner:

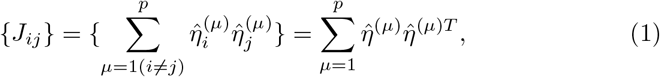

where subscripts *i* and *j* indicate elements (or components) of vector 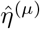. The brackets {…} enclose elements of a matrix shown below. The index *μ* designates the different stored patterns. The entity summed over in the last part of the equation, for *μ* = 1 for instance, is a matrix obtained mathematically by multiplying the column vector 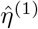 with the corresponding row vector 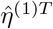, its transpose.

We will first illustrate how 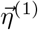 and 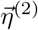 are stored in their normalised forms namely, 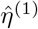 and 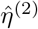, and then describe how normalisation could take place in the EIN-EIN network.

For *μ* = 1, the matrix multiplication as in eqn.(1) is represented in terms of elements as,

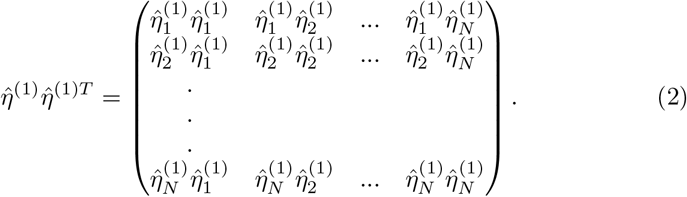

As stated above the elements of this matrix are stored in the SiIN dendrites and for this we need to explore how the multiplicative interactions between the various EIN inputs to the SiINs happen. The axons of EINs in group B each carry one component. For example, 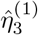 and 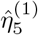 are carried to an SiIN-dendrite by two EIN axons, which together make a synaptic contact with an SiIN-dendrite, and the product of the two inputs 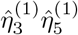 modifies the synapse. This multiplication may reflect particular features of EIN to SiIN synapses, which exhibit short-term facilitation rather than the short-term depression seen for other EIN synapses (Jia and Parker 2016). Facilitation evokes supralinear responses over spike trains (Parker 2003b), and thus could perform a multiplication-like operation of EIN inputs by increasing the gain of SiIN spiking, indicative of a multiplicative effect (Koch and Poggio 1992; Silver 2010).

This matrix multiplication sets up the first stored pattern in the SiIN pool as 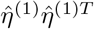 using the othonormalised form of the incident raw vector 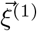. When a second vector, 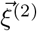 comes in, a set of descending commands goes to SiINs and to the group A EINs. The descending input 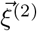 that goes directly to the dendrites of the SiINs passes through the synapses that were modified by 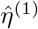, received from the EINs after being processed (i.e. normalised). Thus, the vector 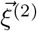 enters SiINs weighted with matrix {*J*_*ij*_} or 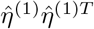. That is, the new descending input is multiplied by 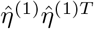, the factor by which the EIN-SiIN synapses were modified after the first input pattern was stored as a postsynaptic change in the SiIN dendrites. The result of this multiplication namely 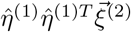, which is a vector, causes a certain magnitude of spiking in the SiINs, and this is sent as an inhibitory input (i.e. with a negative sign), element by element, through monosynaptic SiIN connections to the group A EINs (Parker 2003b). The elements or components of this vector are constructed as explained below.

Consider, as an example, the third row in the matrix of eqn.(2) — it will be multiplied, element by element, with the elements of the column vector 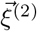 on various contact points on the dendrites of an SiIN and summed by the SiIN like this: 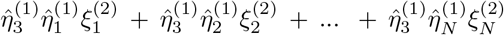. This will constitute one element of a column vector, 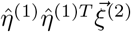. The *N* elements (or components) constructed in this manner will cause spiking of the SiINs that will result in an inhibitory input to the EINs, which, in turn, will get subtracted, element by element, from the vector 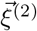 that has also been received by the EINs in group A. In this way the EINs combine two inputs – the inhibitory one from the SiINs and the descending commands from the brain – to construct 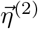 as in eqn.(3) below,

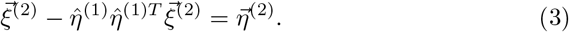

The diagonal elements of the matrix in eqn(2) are set to zero to represent the lack of synapses made by an EIN onto itself.

The resulting 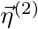 is normalised in the group B EIN network to prepare 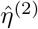 by the following scheme: The 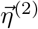 constructed on the network of group A EINs will evoke activity (i.e. a pattern of firing and quiescence) on the group B EINs. The intensity (or magnitude) of activity on each cell, which represents the components (or elements) of 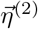, will be different, and a quasi-stable pattern of firing and quiescent cells will emerge. The measure of this collective activity integrated over all the cells in group B can be the magnitude (or length, or norm) of the vector 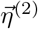, i.e. 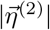.

In contrast to the EIN inputs to the SiINs, the connections between the EINs typically undergo short-term depression. Excitation of the group B EIN pool mediated by synapses with short-term depression can in principle allow division of the activity in each EIN cell by the norm or the length of 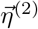, i.e. 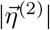, and lead to normalisation of the vector (see Rothman et al 2009 for an example). Thus the normalised 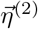, represented by 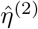 has its *i*^*th*^ component as,

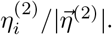

The 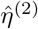, thus generated will comprise +*/*−[fractions] rather than +*/*−1, which when squared and summed will yield 1. That is, the cells will fire or remain quiescent in varying degrees. It is then relayed to the group B EIN-SiIN synapses where it is stored in a Hebbian manner as outlined above in the SiIN dendrites together with 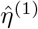 as 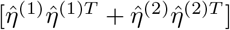.

Note that the eqn.(3) essentially tells the EINs how to compute the differences between 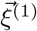 and 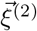, which constitute the 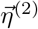. These are processed in the group B EIN-network to prepare the normalised 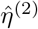, which is then stored in group B EIN-SiIN synapses. Subsequent inputs build on this input to form a library of motor patterns. Every new descending input passes through the same synapses that the first input went through, and the output resulting from the computations outlined above are stored in the SiINs. For example, when 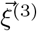 comes in, it is multiplied by 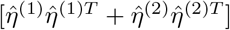 in the dendrites of SiINs, and this product generates an inhibitory output from the SiINs that goes to EINs where it is subtracted from 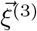, which is received in parallel by the EINs to give 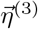,

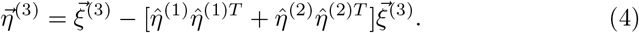

This is then normalised in the group B EIN network and the normalised version, 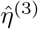, is relayed for storage in the SiIN dendites as outlined above. Thus, the first pattern stored in the SiIN dendrites as 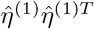 serves as a base to which the 2nd input is attached and stored; 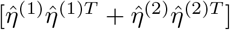 helps in the formation and storage of 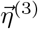; and further on in the sequence, 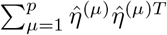, which is the sum total of *p* stored patterns, provides a platform on which 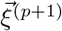 could arrive and convert into 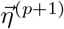 as,

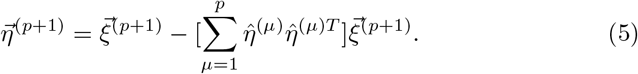

The above scheme outlines how a series of motor commands from the brain could be stored in the spinal cord as schema through identified feedforward and feedback interactions between the excitatory EINs and the inhibitory SiINs. This is illustrated in Fig.1(a) for the very ‘first’ pattern when the system is believed to be a *tabula rasa*, and in Fig.1(b) after the system has stored *p* patterns.

**Figure 1:**
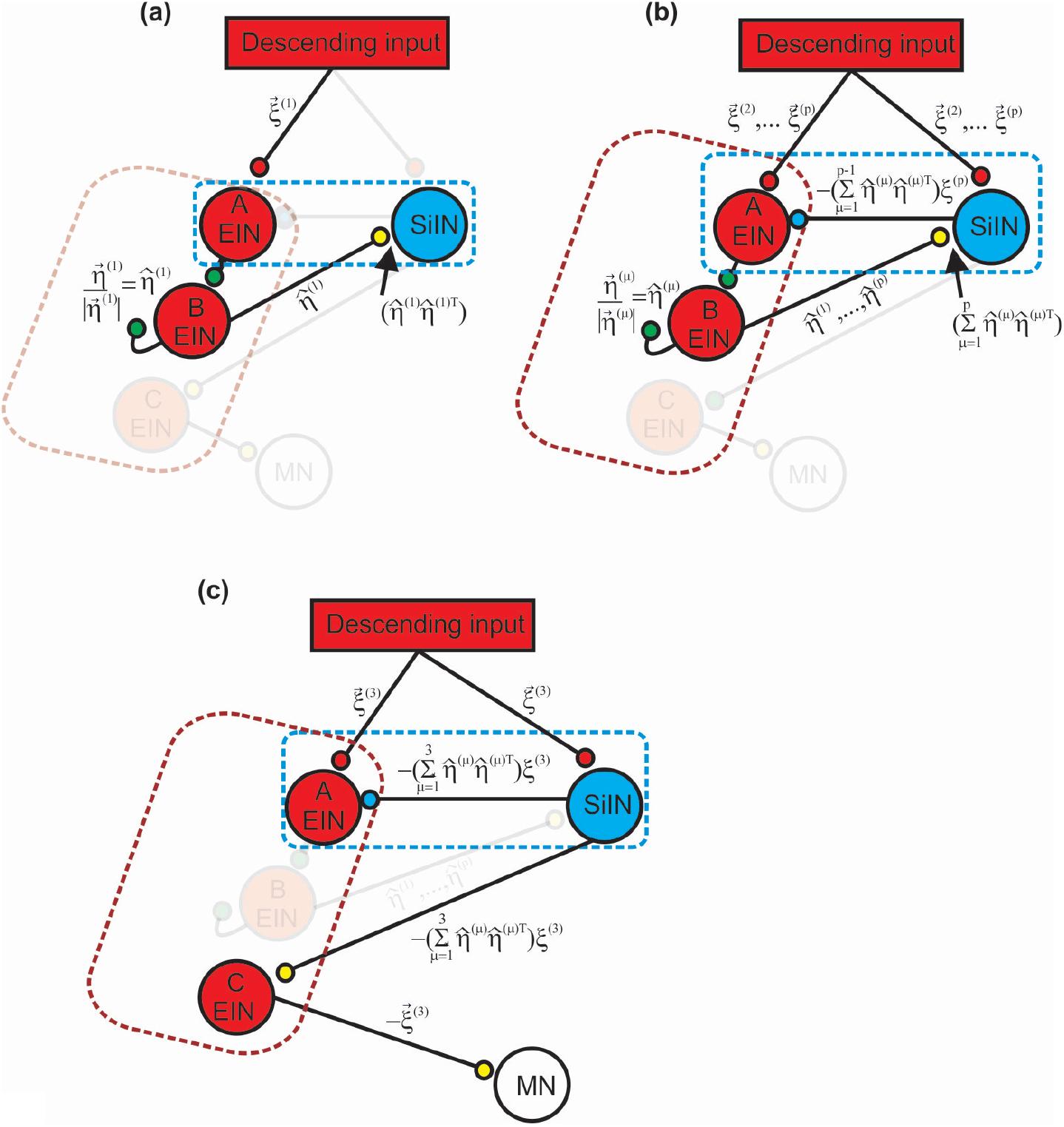
Schematic representation of the circuitry for storage, (a) and (b), and retrieval, (c), of instructions of motor actions as they descend from the brain: (a) shows storage of the very first instruction (or vector) when the system is assumed to be naive, or in *tabula rasa*, and (b) shows storage of the subsequent instructions; (c) demonstrates how an arbitrary instruction descending from the brain is extracted from the library of all the stored instructions that reside in the EIN-SiIN synapses and transmitted to the motor neurons.

## 3. Retrieval from memory

While the above scheme outlines how different motor programs can be stored by building on each other, it is necessary to address how a stored motor program can be retrieved by a particular descending input. While much effort has focused on storage in memory systems (e.g. long-term potentiation) less attention has been paid to how a specific memory can be retrieved.

Let us say that a goal-directed command in a given situation results in 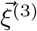 being sent from the brain into the spinal cord. It will go to the dendrites of SiINs and also to those of the group A EINs. It will be received in SiINs with the weight, 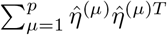 as described after eqn.(4). Since 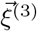 is orthogonal to all 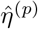’s with *p* > 3, the weighted descending input, which is the post synaptic potential (psp) on SiINs, will simplify to the following reduced form and depolarise the SiINs,

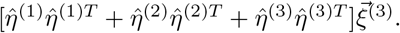

This will go to the group A EINs as an inhibitory input from SiINs (i.e. with a negative sign). Note that mathematically this simplifies to 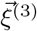 (Srivastava et al., 2000, 2008, 2014), so essentially what will be received by the group A EINs is 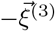. The group A EINs also receive 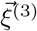 descending from the brain, where it will combine with 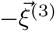 coming from the SiINs and the two inputs will neutralise each other. That 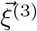 and 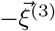 arriving respectively from the brain and SiINs neutralise each other on the group A EINs ensures that nothing is sent back from EINs to SiINs when a previously stored descending command is received, and as a result the stored motor commands are not disturbed.

We suggest that the SiINs will simultaneously transmit the processed output, namely 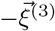, as a command to the group C EINs. This inhibitory input drives the activity in these EINs through a rebound mechanism, and this is relayed onto the MNs to generate the specific motor output associated with the particular descending command (see Discussion for a justification of this approach).

An example demonstrates the retrieval quantitatively in Table 1 (above) while the entire process is illustrated in Figure 1(c) (below).

**Table 1:**
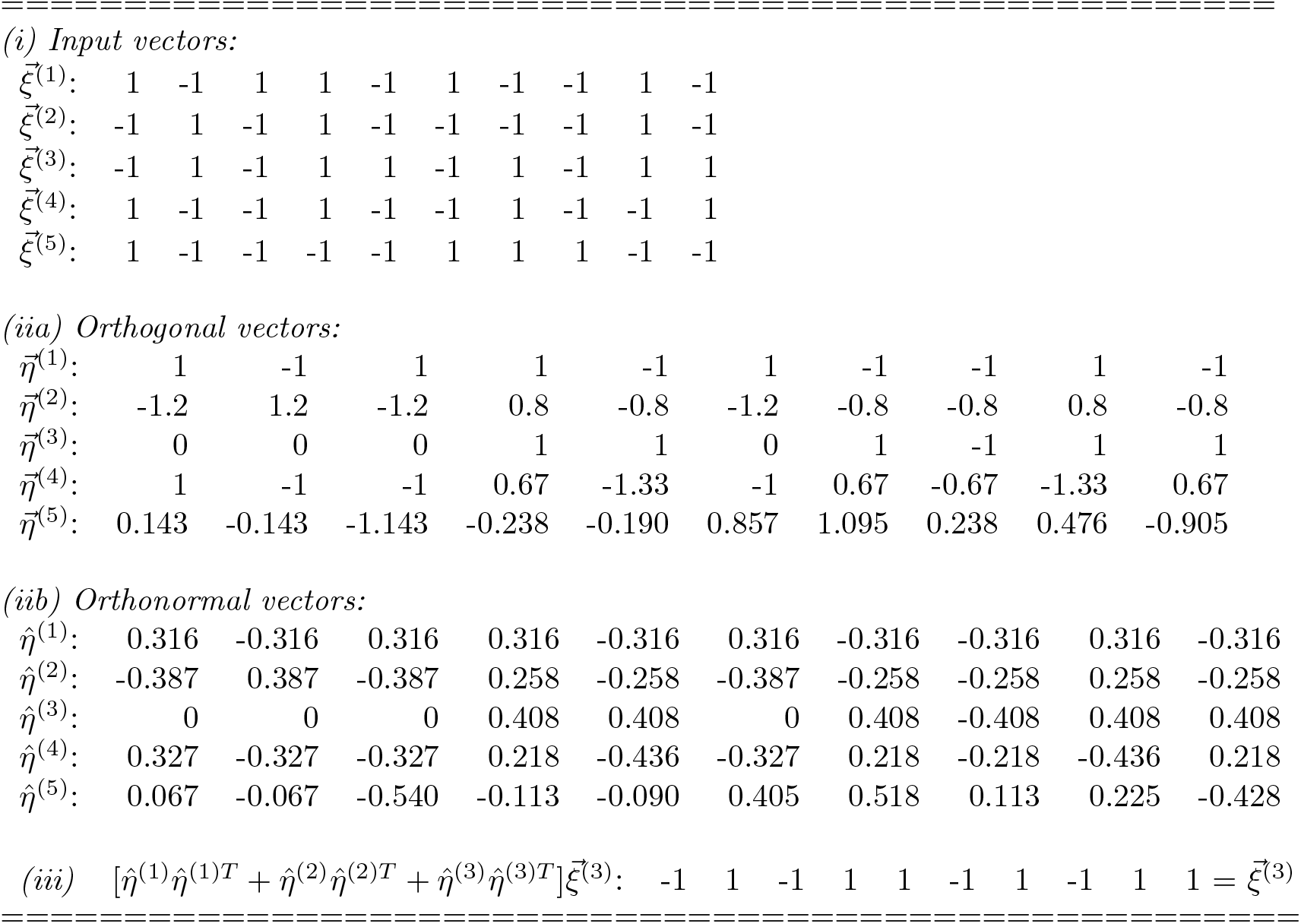
An example showing (i) five patterns representing instructions/signals for motor actions, (iia) their orthogonalized, and (iib) orthonormalized forms that are learned and stored in the spinal cord, and (iii) how, when sent down again by the brain, one of these signals, 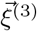, can be reconstructed on SiINs and transmitted to motor neurons via group C EINs to trigger the desired action.

## 4. Discussion

We have suggested a mechanism for how the spinal cord can store motor commands that are encoded and triggered by descending inputs from the brain (Flash and Bizzi 2016). The spinal cord can generate different outputs in response to differing descending inputs (Pierrot-Deseilligny and Burke 2005; see Zelenin et al. 2001), but it is not known how these different responses are programmed into spinal cord locomotor circuits. While the spinal cord was not traditionally considered a site of motor memories, this view has changed (Wolpaw and Tennissen 2001). The mechanisms we suggest are consistent with known circuitry and synaptic properties in the lamprey (and other) spinal cords. The proposed mathematical schemes for orthogonalisation and normalisation require that a number of aspects are tested. These are outlined below.

We suggest that the inhibitory SiINs are the principal site for the storage and retrieval of stored patterns. After the first input has been stored, subsequent inputs come to SiIN dendrites where they are weighted (i.e. multiplied) with the matrix stored in EIN-SiIN synapses. The result is sent as an inhibitory input from SiIN to the group A EINs, where it is subtracted from the descending input at the EINs to produce an orthogonalised vector. This is sent to the group B EINs where it is normalised and sent to the SiIN dendrites to be added to the previously stored ortho-normalised vectors (in a matrix form). The main point about storage is that the raw input vectors are stored in EIN-SiIN synapses in a Hebbian manner in the form of a matrix whose elements are sums of the components of the ortho-normalised vectors constructed from the raw input vectors.

For retrieval, e.g. of 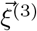, a stored program needs to be reconstructed via the SiINs. We suggest that subsequent to its storage the first time, 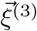 attaches with a convolution of all the stored motor actions (which may be say, *p* in number) in their orthonormalised form; the convolution in mathematical terms is: 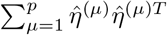 and resides in synapses made by EINs in the SiIN dendrites. This attachment causes all the motor actions to be filtered out on retrieval of 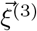 except the first three since 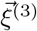 is orthogonal to (or, has no overlap with) all 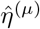’s with *μ* > 3. Thus, 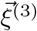 connects as 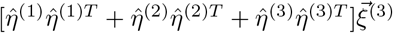. This results in reconstruction of 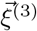 on the SiINs, which send 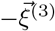 back to the group A EINs where it will add with 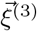 to cancel out the descending command to the group A EINs; this ensures that the group B EINs are not activated, and will not re-consolidate the descending command when it is retrieved (note, however, that re-consolidation is not without precedent in memory systems; McKenzie and Eichenbaum 2011).

The central role for inhibition that we propose contrasts the focus on excitatory interactions in driving motor outputs (Kiehn 2010), and for memory storage generally (Bliss and Collingridge 2013). Inhibitory interneurons like the SiINs are common in spinal cord networks (Roberts et al 2010; Higashijim et al. 2004; Kiehn et al. 2010). They have been removed from one scheme for the lamprey locomotor circuit (Grillner 2003), but not other schemes (see Buchanan 2001). The removal was prompted by the failure to simulate the effects of 5-HT on the locomotor network when this inhibition was present (Hellgren et al. 1992), and is thus an idealisation to allow the model to match the biological system (see Parker 2006). On its own this doesn’t negate a role for the SiINs and the theoretical scheme we propose here.

It could be argued that the scheme we propose is negated because fictive locomotor activity persists when inhibition is blocked (see, for example, Alford et al. 1990). However, this persistence of activity with excitation alone does not show that inhibition is unimportant. Firstly, fictive activity cannot be assumed to directly reflect normal outputs (see Jackson et al. 2005; Messina et al 2017; Parker and Srivastava 2013); and secondly, removing inhibition has diverse effects on motor outputs: it alters the frequency, causes synchronous activity across the two sides of the body, or disrupts the motor output (Alford et al. 1990; Cohen and Harris-Warrick 1984; Gosgnach et al., 2006; Hagevik and McClellan 1994; McPherson et al., 1994; Roberts et al. 1985). Berg et al (2007) suggest that instead of a model that assumes separate phases of inhibition and excitation, locomotor activity reflects a balance between inhibition and excitation, consistent with a more involved role for inhibition in motor outputs.

We suggest that the 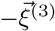 from the SiINs leads to the excitation that evokes the retrieved motor activity. While action potentials are conventionally evoked by excitatory inputs, phasic inhibition can drive activity by evoking rebound firing (post-inhibitory rebound or PIR). PIR is defined as depolarization evoked by decay of hyperpolarization (which here will come from the SiINs). There is evidence that this drives excitation in diverse systems. In the superior paraolivary nucleus of mice, the classical roles for excitation and inhibition are switched, with inhibitory inputs driving spike firing via a PIR mechanism and excitatory inputs modulating this response (Felix et al 2011). A similar reliance on rebound excitation from inhibition rather than excitation is seen in the rat entorhinal cortex (Ferrante et al 2017), where it contributes to the storage of spatial memories (Pastoll et al 2013). PIR contributes to locomotor rhythms in the spinal cord of the rat (Bertrand and Cazalets 1998), lamprey (Matsushima et al. 1993) and xenopus (Roberts et al. 2010), and in motor activity in a range of invertebrates (Arshavsky et al. 1998; Straub and Benjamin 2001; Fox et al 2013; Angstadt et al 2005). PIR is thus a common mechanism for evoking excitation.

A crucial, and testable, aspect of the proposed storage mechanism is that the SiINs can generate supralinear responses to paired EIN inputs. Multiplicative effects occur in the nervous system (see Silver 2010 for review). While, multiplicative mechanisms are not well understood, they could reflect non-linear synaptic interactions or the activation of voltage-activated dendritic conductances (see Silver 2010). The SiINs are of interest in this respect, as unlike most synapses in the lamprey locomotor network, the synapses they receive from the EINs facilitate rather than depress (Jia and Parker 2016; Parker 2003b). While the functional role of short-term plasticity in spinal and other neural circuits is still unclear (Anwar et al. 2017; Bertrand and Cazalet 2013; Jia and Parker 2016), facilitation may underlie the non-linear dendritic interactions underlying the matrix multiplication needed for storage of motor patterns in the SiINs 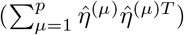 (Silver 2010). Conversely, depression of EIN-EIN connections could provide a divisive mechanism (Silver 2010) that could allow normalisation of 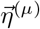 to 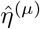.

The temporal relationship of descending inputs to the EINs and SiINs also needs to be considered. The storage of the initial pattern does not involve direct activation of the SiINs, only storage of effects in their dendrites after activation of the EINs. However, the SiINs have to be activated by the 2nd (and subsequent) descending input before the EINs. This is in principle possible by the specific projections of the descending input pathways (Lemon 2008).

This scheme thus raises a number of testable hypotheses. Examining these requires direct control over the descending pathways in an intact, behaving, and initially naïve system on which imposed patterns can be manipulated at different stages in the appropriate temporal pattern. This may be possible in a simpler genetically tractable system like the zebrafish where the activity of different classes of neurons could be controlled during early development. For example, the activity of the appropriate inhibitory interneurons (e.g. engrailed-1 positive interneurons; Higashijima et al. 2004) could be regulated using optogenetics. Manipulating these cells should affect motor performance if these interneurons have a general role in motor activity. However, it should be possible to determine the effects of removing these neurons on basic locomotor behaviours, and if this is not too severe this could allow comparison of this deficit to the failure to learn or retrieve motor responses to a particular stimulus. It may also be possible to provide specific descending commands that could be stored as motor memories in the way we propose, the initial pattern through the EINs, and subsequent patterns through the SiINs and then the EINs, thus allowing the role of inhibition in storage to be tested. The capacity of a system without orthogonalisation is 0.14N (where N is the number of neurons), but is close to N when orthogonalised inputs are stored in the way we suggest here (Srivastava et al. 2008, 2014). This predicts that reducing the inhibitory cell population will result in a reduced number of stored patterns.

Learning of motor commands could also reflect recovery of function after spinal cord injury. A complete lesion will remove all descending commands, isolating sub-lesion networks and preventing goal-directed motor outputs. Incomplete lesions could either reflect spared axons, or regeneration of damaged axons. Regeneration occurs spontaneously in lower vertebrates like the lamprey, but not in mammals where promoting regeneration remains the dominant focus of research into spinal cord injury (Steward et al. 2012). In either case, the descending command, and thus the resulting activation vector 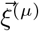, will be disrupted. For effective functional recovery the descending input will have to be re-learnt. Complete locomotor recovery occurs spontaneously in the lamprey. While this has been considered to reflect regeneration (Cohen et al 1988), there are also changes in the spinal cord above and below the lesion site that suggest additional recovery factors (Parker 2017), and the need to consider how regenerated inputs interact with the sub-lesion spinal cord to generate the same output using different circuitry with different properties (see Hoffman and Parker 2011). There is an increase in inhibitory activity below, but not above the lesion site.

This may relate to the dependence on inhibition in the model that we propose here. However, this will also need to be tested.

## Acknowledgements

VS would like to thank the University of Hyderabad for hospitality during the course of this work. We are grateful to Dr A Ramesh Naidu for help in the computations in Table 1.

## Notes

### Competing Interest Statement

The authors have declared no competing interest.

## References

Alford S, Sigvardt KA, Williams TL (1990) GABAergic control of rhythmic activity in the presence of strychnine in the lamprey spinal cord. Brain Research 506:303–306.

Ampatzis K, Song J, Ausborn J, Manira AE. 2014. Separate Microcircuit Modules of Distinct V2a Interneurons and Motoneurons Control the Speed of Locomotion. Neuron 83: 934–43.

Angstadt JD, Grassmann JL, Theriault KM, Levasseur SM. 2005. Mechanisms of postinhibitory rebound and its modulation by serotonin in excitatory swim motor neurons of the medicinal leech. Journal of Comparative Physiology A 191: 715–32.

Anwar H, Li X, Bucher D, Nadim F. 2017. Functional roles of short-term synaptic plasticity with an emphasis on inhibition LA - eng. Current opinion in neurobiology 43 AN - 28122326: 71–8.

Arshavsky Y, Orlovsky G, Panchin Y, Roberts A, Soffe S. 1993. Neuronal control of swimming locomotion: analysis of the pteropod mollusc clione and embryos of the amphibian Xenopus. TINS 16: 227–33.

Ausborn J, Shevtsova NA, Caggiano V, Danner SM, Rybak IA (2019) Computational modeling of brainstem circuits controlling locomotor frequency and gait. eLife 8:e43587.

Ayers J, Carpenter G, Currie S, Kinch J. 1983. Which behavior does the lamprey central motor program mediate? Science 221: 1312–4.

Berg R, Alaburda A, Hounsgaard J (2007) Balanced inhibition and excitation drive spike activity in spinal half-centers. Science 315:390–393.

Bertrand S Cazalet J-R (2013) Activity-dependent Synaptic Plasticity and Metaplasticity in Spinal Motor Networks. Current Pharmaceutical Design 19: 4498–508.

Bizzi E, Cheung V, d’Avella A, Saltiel P, Tresch M (2008) Combining modules for movement. Brain Res Rev 57:125–133.

Bliss TV, Collingridge GL (2013) Expression of NMDA receptor-dependent LTP in the hippocampus: bridging the divide. Molecular Brain 6:5.

Brodin L, Grillner S, Rovainen C. 1985. N-methyl-D-aspartate (NMDA), kainite and quisqualate receptors and the generation of fictive locomotion in the lamprey spinal cord. Brain Res 325: 302–6.

Buchanan J (2001) Contributions of identifiable neurons and neuron classes to lamprey vertebrate neurobiology. Prog Neurobiol 63:441–466.

Carandini M, Heeger DJ. 2011. Normalization as a canonical neural computation. Nature Reviews Neuroscience 13: 51–62.

Cohen A, Mackler S, Selzer M (1988) Behavioral recovery following spinal transections: functional regeneration in the lamprey CNS. Trends in Neurosci 11:227–231.

Cazalets J, Sqalli-Houssaini Y, Clarac F. 1992. Activation of the central pattern generators for locomotion by serotonin and excitatory amino acids in neonatal rat. J Physiol 455: 187–204.

Cohen AH, Harris-Warrick RM. 1984. Strychnine eliminates alternating motor output during fictive locomotion in the lamprey. Brain Research 293: 164–7.

Deliagina T, Zelenin P, Orlovsky G (2002) Encoding and decoding of reticulospinal commands. Brain Res Rev 40:166–177.

Felix RA, 2nd, Fridberger A, Leijon S, Berrebi AS, Magnusson AK. 2011. Sound rhythms are encoded by postinhibitory rebound spiking in the superior parao-livary nucleus. The Journal of Neuroscience 31: 12566–78.

Flash T, Bizzi E (2016) Cortical circuits and modules in movement generation: experiments and theories. Current Opinion in Neurobiology Microcircuit computation and evolution 41:174–178.

Fox DM, Rotstein HG, Nadim F. 2013. Bursting in Neurons and Small Networks. In Encyclopedia of Computational Neuroscience, ed. D Jaeger, R Jung, pp. 1–17. New York, NY: Springer New York.

French R. 1999. Catastrophic forgetting in connectionist networks. Trends Cogn Sci. 3: 128–35.

Goulding M (2009) Circuits controlling vertebrate locomotion: moving in a new direction. Nat Rev Neurosci 10:507–518.

Gosgnach S, Lanuza GM, Butt SJB, Saueressig H, Zhang Y, et al. 2006. V1 spinal neurons regulate the speed of vertebrate locomotor outputs. 440: 215–9.

Graziano M (2006) The organization of behavioural repertoires in the motor cortex. Ann Rev Neurosci 29:105–134.

Grillner S (2003) The motor infrastructure: from ion channel to neuronal networks. Nat Rev Neurosci 4:573–586.

Hagevik A, McClellan AD. 1994. Coupling of spinal locomotor networks in larval lamprey revealed by receptor blockers for inhibitory amino acids: neurophysiology and computer modeling. Journal of Neurophysiology 72: 1810–29.

Hellgren J, Grillner S, Lansner A (1992) Computer simulation of the segmental neural network generating locomotion in lamprey by using populations of netwrok interneurons. Biol Cybern 68:1–13.

Higashijima S-i, Masino MA, Mandel G, Fetcho JR (2004) Engrailed-1 Expression Marks a Primitive Class of Inhibitory Spinal Interneuron. The Journal of Neuroscience 24:5827–5839.

Hoffman N, Parker D (2011) Interactive and individual effects of sensory potentiation and region-specific changes in excitability after spinal cord injury. Neuroscience 199:563–576.

Jackson A, Horinek D, Boyd M, McClellan A (2005) Disruption of left-right reciprocal coupling in the spinal cord of larval lamprey abolishes brain-initiated locomotor activity. J Neurophsyiol 94(3):2031–2044.

Jia Y, Parker D (2016) Short-Term Synaptic Plasticity at Interneuronal Synapses Could Sculpt Rhythmic Motor Patterns. Frontiers in Neural Circuits http://dx.doi.org/10.3389/fncir.2016.00004.

Kiehn O, Dougherty K, Hägglund M, Borgius L, Talpalar A, Restrepo C (2010) Probing spinal circuits controlling walking in mammals. Biochem Biophys Res Commun 396:11–18.

Koch, C., and Poggio, T. Multiplying with synapses and neurons. Single Neuron Computation (1992), 315–345.

Lemon RN. 2008. Descending Pathways in Motor Control. 10.1146/annurev.neuro.31.060407.125547. Annual Review of Neuroscience 31: 195–218.

Li W-C, Moult PR. 2012. The control of locomotor frequency by excitation and inhibition. The Journal of Neuroscience 32: 6220–30.

Li W, Soffe S, Wolf E, Roberts A (2006) Persistent responses to brief stimuli: feedback excitation among brainstem neurons. J Neurosci 26:4026–4035.

Li W, Roberts A, Soffe S (2009) Locomotor rhythm maintenance: electrical coupling among premotor excitatory interneurons in the brainstem and spinal cord of young Xenopus tadpoles. J Physiol 587:1677–1693.

Matsushima T, Tegner J, Hill R, Grillner S. 1993. GABAB receptor activation causes a depression of low- and high-voltage-activated Ca2+ currents, postinhibitory rebound, and postspike afterhyperpolarization in lamprey neurons. J Neurophsyiol 70: 2606–19.

McKenzie S, Eichenbaum H (2011) Consolidation and reconsolidation: Two lives of memories? Neuron 71: 224–233.

McPherson D, Buchanan J, Kasicki S. 1994. Effects of strychnine on ficitive swimming in the lamprey: evidence for glycinergic inhibition, discrepancies with model predictions, and novel modulatory rhythms. J. Comp. Physiol. A 175: 311–21.

Messina JA SPA, Hargis S, Thompson WE, McClellan AD. (2017) Elimination of Left-Right Reciprocal Coupling in the Adult Lamprey Spinal Cord Abolishes the Generation of Locomotor Activity. Frontiers in Neural Circuits 11: 89.

Parker D (2003a) Variable Properties in a Single Class of Excitatory Spinal Synapse. J Neurosci 23: 3154–3163.

Parker D (2003b) Activity-Dependent Feedforward Inhibition Modulates Synaptic Transmission in a Spinal Locomotor Network. J Neurosci 23: 11085–11093.

Parker D (2006) Complexities and uncertainties of neuronal network function. Philosophical Transactions of the Royal Society B: Biological Sciences 361: 81–99.

Parker D (2010) Neuronal network analyses: premises, promises and uncertainties. Phil Trans R Soc Lond B 365: 2315–2328.

Parker D. 2015. Synaptic Variability Introduces State-Dependent Modulation of Excitatory Spinal Cord Synapses. Neural Plasticity 2015: Article ID 512156.

Parker D (2017) The Lesioned Spinal Cord Is a “New” Spinal Cord: Evidence from Functional Changes after Spinal Injury in Lamprey. Frontiers in Neural Circuits 11: 84.

Parker D, Srivastava V. 2013. Dynamic systems approaches and levels of analysis in the nervous system. Frontiers in Physiology 4: Article 15.

Pierrot-Deseilligny E, Burke D (2005) The circuitry of the human spinal cord: its role in motor control and movement disorders. Cambridge: Cambridge University Press.

Roberts A, Li W, Soffe S (2010) How neurons generate behavior in a hatchling amphibian tadpole: an outline. Front Behav Neurosci 4: 16.

Roberts A, Dale N, Evoy WH, Soffe SR. 1985. Synaptic potentials in motoneurons during fictive swimming in spinal Xenopus embryos. Journal of Neurophysiology 54: 1–10.

Rothman J, Cathala L, Steuber V, Silver R (2009) Synaptic depression enables neuronal gain control. Nature 457: 1015–1019.

Schwartz AB (2016) Movement: How the Brain Communicates with the World. Cell 164: 1122–1135.

Shik ML, Orlovsky GN. 1976. Neurophysiology of locomotor automatism. Physiological Reviews 56: 465–501.

Silver R (2010) Neuronal arithmetic. Nature Rev Neurosci 11: 474–489.

Soffe SR, Roberts A, Li W-C. 2009. Defining the excitatory neurons that drive the locomotor rhythm in a simple vertebrate: insights into the origin of reticulospinal control LA - eng. The Journal of physiology 587: 4829–44.

Srivastava V, Edwards S (2000) A model of how the brain discriminates and categorises. Physica A 276: 352–358.

Srivastava V, Parker D, Edwards S (2008) The nervous system might ‘orthogonalize’ to discriminate. J Theor Biol 253: 514–517.

Srivastava V, Sampath S, Parker D (2014) Overcoming Catastrophic Interference in Connectionist Networks Using Gram-Schmidt Orthogonalization. PlosONE 9(9): e105619.

Srivastava V, Sampath S (2018) Could the brain function mathematically. Neurology and Neuroscience Research 1(1): 1–4.

Steward O, Popovich PG, Dietrich WD, Kleitman N (2012) Replication and reproducibility in spinal cord injury research. Experimental Neurology Special Issue: NIH Replication Studies 233: 597–605.

Straub VA, Benjamin PR. 2001. Extrinsic modulation and motor pattern generation in a feeding network: a cellular study. The Journal of Neuroscience 21: 1767–78.

Talpalar AE, Kiehn O. 2010. Glutamatergic mechanisms for speed control and network operation in the rodent locomotor CpG. Frontiers in neural circuits 4 AN - 20844601: 19.

Thompson E (2009) Habituation: A History. Neurobiol Learn Mem 92: 127–134.

Thompson A, Wolpaw J. 2014. Operant conditioning of spinal reflexes: from basic science to clinical therapy. Front. Integr. Neurosci. 8: Article 25.

Wang H, Jung R. 2002. Variability analyses suggest that supraspino–spinal interactions provide dynamic stability in motor control. Brain Res 930: 83–100.

Wolpaw J, Tennissen A (2001) Activity-dependent spinal cord plasticity in health and disease. Ann Rev Neurosci 24:807–843.

Zelenin PV, Grillner S, Orlovsky GN, Deliagina TG (2001) Heterogeneity of the Population of Command Neurons in the Lamprey. J Neurosci 21:7793–7803.

